# Profiling the surface proteome identifies actionable biology for TSC1 mutant cells beyond mTORC1 signaling

**DOI:** 10.1101/382929

**Authors:** Junnian Wei, Kevin K. Leung, Charles Truillet, Davide Ruggero, James A. Wells, Michael J. Evans

## Abstract

Loss of the TSC1/TSC2 complex leads to constitutively high mTORC1 signaling; however, pharmacological inhibition of mTORC1 in this setting produces a broad spectrum of clinical responses. We report herein several cell surface proteins upregulated by inactivation of TSC1 that present therapeutic alternatives or adjuvants to direct mTORC1 inhibition. A proteomics screen revealed that TSC1 loss most dramatically induced the expression of neprilysin (NEP/CD10) and aminopeptidase N (APN/CD13). The survival of TSC1 null human cancer cells was dependent on NEP expression, and TSC1 mutation sensitized cells to biochemical inhibition of APN. Remarkably, NEP and APN upregulation occurred via a TSC2- and mTORC1-independent mechanism; therefore, the antiproliferative effects of mTORC1 inhibition could be augmented by co-suppression of APN activity.

**Statement of significance:** These data introduce a non-canonical biological role for TSC1 beyond regulating mTORC1 signaling, which also enabled several immediately translatable therapeutic strategies for clinically problematic cells with TSC1 mutations.

## Introduction

Somatic or germline genetic mutations that inactivate TSC1 or TSC2 contribute to the pathogenesis of several deadly cancers (e.g. bladder, kidney) and debilitating human disorders (e.g. tuberous sclerosis complex, focal cortical dysplasia)^1^. Under normal conditions, TSC1 is known to heterodimerize with TSC2 to provide protection from ubiquitin mediated degradation^2^, while TSC2 employs a GTPase activating protein domain to biochemically convert GTP-Rheb to GDP-Rheb^3^. As GTP-Rheb is required for the activation of mTORC1, loss of the TSC1/TSC2 complex generally results in constitutively high mTORC1 signaling. On this basis, there has been considerable interest in treating clinically problematic cells with inactivating mutations in TSC1 and/or TSC2 by inhibiting mTORC1 signaling^4^.

While TSC1/TSC2 null preclinical models are sensitive to mTORC1 inhibitors^5–7^, the clinical experience has been comparatively more perplexing. For instance, benign and malignant tubers arising from tuberous sclerosis complex are highly responsive to the rapalogue everolimus (RAD001)^8–10^. Moreover, somatic TSC1 or TSC2 mutations in bladder, kidney, thyroid, and liver cancers have been associated retrospectively with clinical responses to everolimus, including examples of long term bladder and thyroid cancer remissions^11–15^. However, not all patients with TSC1 and/or TSC2 mutations experience durable radiographic responses, and it is currently unclear if TSC1 and/or TSC2 mutational status bears any predictive value in treatment naïve patients. These observations could suggest that additional pathobiological events are required to confer sensitivity to mTORC1 inhibition, and/or that loss of TSC1/TSC2 imparts pathobiological effects independent of mTORC1 signaling.

Collectively, these considerations motivated us to apply an unbiased molecular profiling approach to discover alternative therapeutic strategies. Protein signaling on the cell surface is a prominent mechanism of intra- and intercellular communication, and an important component of cellular transformation. Moreover, the accessibility of cell surface proteins enables comparatively more and chemically diverse therapeutic strategies than are available for targeting intracellular proteins, including clinically validated technologies like antibody-drug conjugates and radioimmunotherapy^16^. However, to our knowledge, cell surface protein changes associated with loss of TSC1 or TSC2 had not yet been defined. On this basis, we hypothesized that profiling the cell surface proteome in genetically defined cell line models might identify new therapeutic targets arising from TSC1/TSC2 loss.

## Results

We conducted a LC-MS/MS study to define the cell surface proteome in isogenic pairs of *Tsc1*^-/-^ and *Tsc1*^+/+^ mouse embryonic fibroblasts (MEFs). Isogenic pairs of MEFs were selected as the complete genetic knockout of TSC1 would allow for a rigorous binary comparison of proteomic changes due to loss of TSC1. A SILAC quantification scheme was coupled to an N-glycan enrichment approach to isolate and analyze cell surface proteomes^17,18^. Two SILAC experiments performed (light *Tsc1*^-/-^ vs heavy *Tsc1*^+/+^, heavy *Tsc1*^-/-^ vs light *Tsc1*^+/+^) after 6 hours of serum starvation resulted in quantification of 5197 unique peptide spectra corresponding to 558 protein IDs. The enrichment ratio (either heavy:light or light:heavy) for each extracellular protein identified was calculated using R and showed strong reproducibility (**Supplemental Figure 1A–B**). Overall, 20 proteins were predicted to have a median log_2_ enrichment ratio ≥ 2 (*Tsc1*^-/-^ versus *Tsc1*^+/+^) with p < 0.05 (**Figure 1A, Supplemental figure 1C, Table 1**). Among these proteins, we were encouraged to see that the transferrin receptor (*Tfrc*) and glucose transporter 1 (*Slc2a1*) were overexpressed on the surface of *Tsc1*^-/-^ MEFs, consistent with prior literature showing that mTORC1 upregulates their respective cell surface expression (**Figure 1B**)^19–21^. Gene set enrichment analysis (GSEA) of the differentially regulated hits on the cell surface using GO term gene sets identified a significant de-enrichment of genes sets annotated with biological adhesion (**Supplemental Figure 2A–D**). Further GSEA using a combination of KEGG, REACTOME, BIOCARTA, and HALLMARK gene sets identified a de-enrichment of the focal adhesion pathway (**Supplemental Figure 2E–G**). The specific proteins identified in the leading edge of the de-enriched KEGG focal adhesion pathway included *Itga9*, *Itgb3*, *Pdgfra*, *Flt4*, *Egfr*, *Ctnnb1*, *Itga1*, *Erbb3*, *Itga6*, *Mst1r*, and *Itga3* (**Supplemental Figure 2H**). This finding is consistent with previous literature showing that TSC1 regulates cellular adhesion^22^.

**Figure 1:**
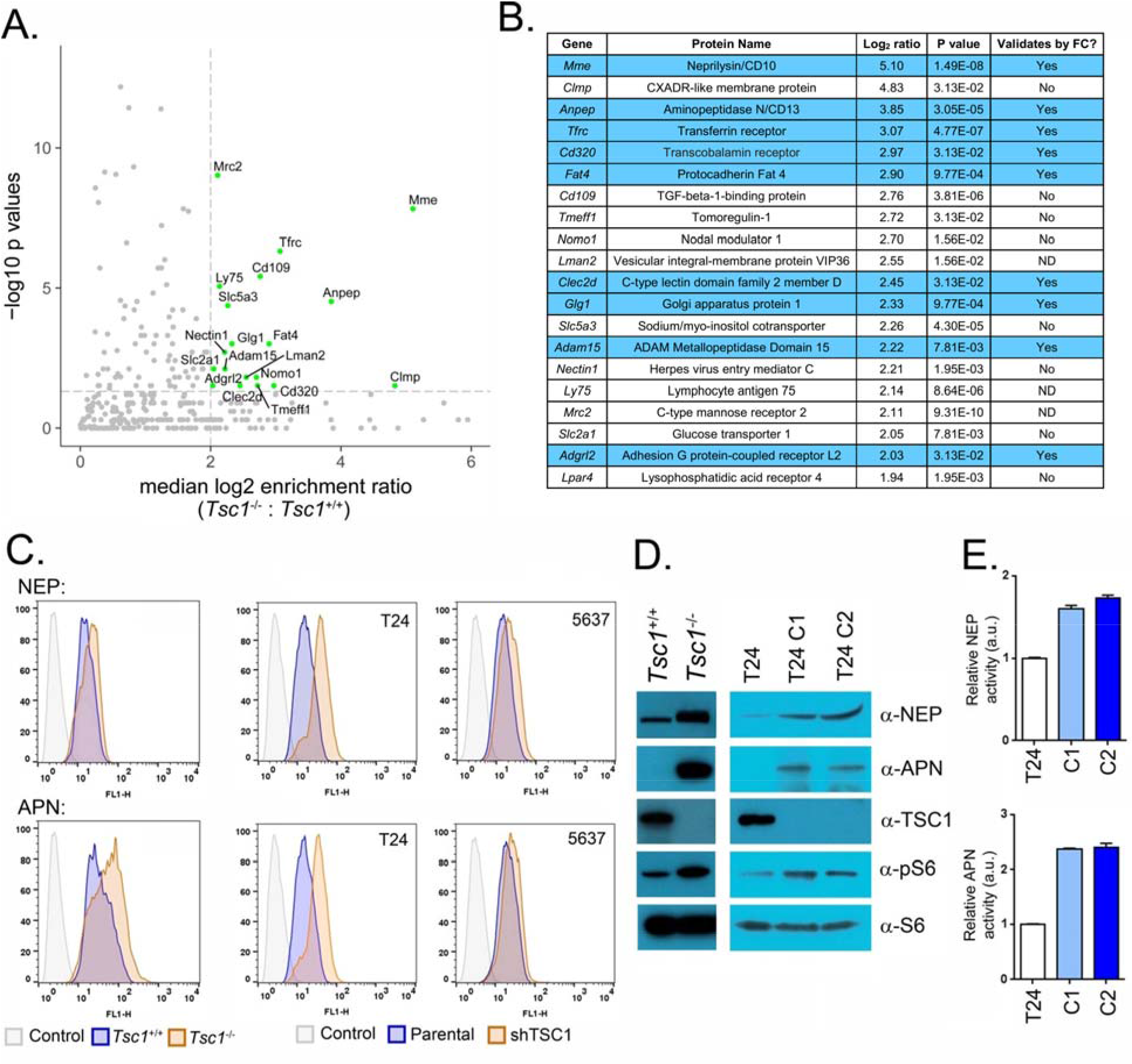
Proteomic profiling and flow cytometry identifies many proteins upregulated on the cell surface in TSC1 null cells.

**A.** A portion of a volcano plot showing the hits detected as upregulated in the cell surface proteome of *Tsc1*^-/-^ versus *Tsc1*^+/+^ MEFs. The hits labeled in green represent those that had a log_2_ fold enrichment > 2, and were statistically significant (P < 0. 05). Those hits shaded in grey represent proteins detect in the proteomics screen that were not significantly altered. **B.** A table summarizing the top 20 hits predicted to be upregulated in *Tsc1*^-/-^ MEFs by LC-MS/MS, and the results of the hand validation by flow cytometry. Hits were determined to validate if the percent change in mean fluorescence intensity between isogenic pairs was > 20%. Hits that validated are shaded in blue, and all hits were validated in at least two independent flow cytometry experiments. **C.** Representative histograms showing that NEP and APN are upregulated in *Tsc1*^+/+^ and *Tsc1*^-/-^ MEFs (left) and T24 and 5637 sublines with TSC1 knockdown by shRNA compared to the parental cell line (right). **D.** Immunoblot data showing the expression differences for NEP and APN in whole cell lysates of *Tsc1*^-/-^ and *Tsc1*^+/+^ MEFs, and parental T24 compared to single cell clones T24 C1 and T24 C2 harboring *Tsc1* knockout via CRISPR/Cas9. **E.** Activity data showing higher levels of NEP and APN biochemistry in T24 C1 and C2 compared to parental T24. Biochemical activity was assay from intact cells using a commercial fluorometric assay for substrate hydrolysis.

Flow cytometry was used to validate the overexpression of the 20 hits predicted to be upregulated in *Tsc1*^-/-^ MEFs by LC-MS/MS (**Figure 1B and Supplemental Figure 3A**). Five of the hits were determined to be upregulated by ≥ 20% (log scale) in the *Tsc1*^-/-^ versus *Tsc1*^+/+^ cell surface proteome: neprilysin (NEP/CD10, *Mme*), aminopeptidase N (APN/CD13, Anpep), the transferrin receptor (TFRC, *Tfrc*), Golgi apparatus protein 1 (GSLG1, *Glg1*), and adhesion G protein coupled receptor L2 (AGRL2, *Adgrl2*, see **Supplemental Figure 3B**). We next tested if the cell surface expression of the two top validated hits, the metalloproteases NEP and APN, was increased by loss of TSC1 in human cancer models representative of malignancies in which TSC1 mutation is common. TSC1 was stably suppressed by shRNA in the human bladder cancer cell lines T24 and 5637 (**Supplemental Figure 4A** and **4B**). Both NEP and APN were upregulated on the surface of the shTSC1 sublines of T24 and 5367 compared to the parental cell line (**Figure 1C**). Probing for NEP and APN expression changes via immunoblot showed that expression levels for each protein were substantially higher in *Tsc1*^-/-^ versus *Tsc1*^+/+^ MEFs (**Figure 1D**). Relative mRNA levels of *Mme* and *Anpep* were also significantly higher in *Tsc1*^-/-^ compared to *Tsc1*^+/+^ MEFs by rtPCR (**Supplemental Figure 5A**). Moreover, analysis of gene expression profiling data previously reported in the literature predicted higher levels of *Mme* and *Anpep* mRNA in *Tsc1*^-/-^ versus *Tsc1*^+/+^ MEFs (**Supplemental Figure 5B**)^23^.

To compare protein expression changes in a human cell line bearing TSC1 knockout, TSC1 was deleted from T24 cell line using CRISPR/Cas9 technology. Two single cell clones (T24 C1, T24 C2) treated with discrete sgRNAs were isolated, expanded and validated for TSC1 knockout (**Figure 1E**). NEP and APN expression levels were significantly higher in the TSC1 knockout sublines compared to parental T24 (**Figure 1E**). NEP and APN biochemical activities were also significantly higher in viable T24 C1 and C2 cells compared to the parental line (**Figure 1F** and **Supplemental Figure 6A**).

We next tested whether NEP and APN expression and/or activity is required for survival and proliferation in a panel of four endogenously TSC1 null human cancer cell lines—the bladder cancer cell lines HCV29 (TSC1 Q55X), RT4 (TSC1 L557fs), 97-1 (TSC1 R692X), and the thyroid cancer cell line 8505 C (TSC1 R692Q, see **Supplemental Figure 6B**). In vitro, genetic suppression of *Mme* or *Anpep* with siRNA potently reduced proliferation in all cell lines compared to treatment with a non-targeting siRNA (**Figure 2A** and **Supplemental Figure 7A** and **7B**). Interestingly, treatment with the biochemical NEP inhibitor LBQ657, the bioactive form of the heart failure drug Sacubitril, did not impact proliferation of the cells within 5 days (**Figure 2B and Supplemental Figure 8**). On the other hand, treatment with the biochemical APN inhibitor CHR2797 resulted in significant antiproliferative effects at 3 and 5 days (**Figure 2B** and **Supplemental Figure 9A**). Treatment with bestatin, a comparatively less potent and specific biochemical inhibitor of APN, did not inhibit proliferation at 20 μM (**Figure 2B** and **Supplemental Figure 9B**). Further analysis of CHR2797 treatment effects showed an induction of caspase 9 biochemical activity in all cell lines, consistent with cell death via apoptosis (**Supplemental Figure 10A**). Moreover, the growth of 97-1 and 8505 C tumors implanted subcutaneously in *nu/nu* mice was suppressed by daily i.p. CHR2797 treatment over 20-26 days compared to tumors implanted in mice exposed to vehicle (**Figure 2C** and **Supplemental Figure 10B**).

**Figure 2:**
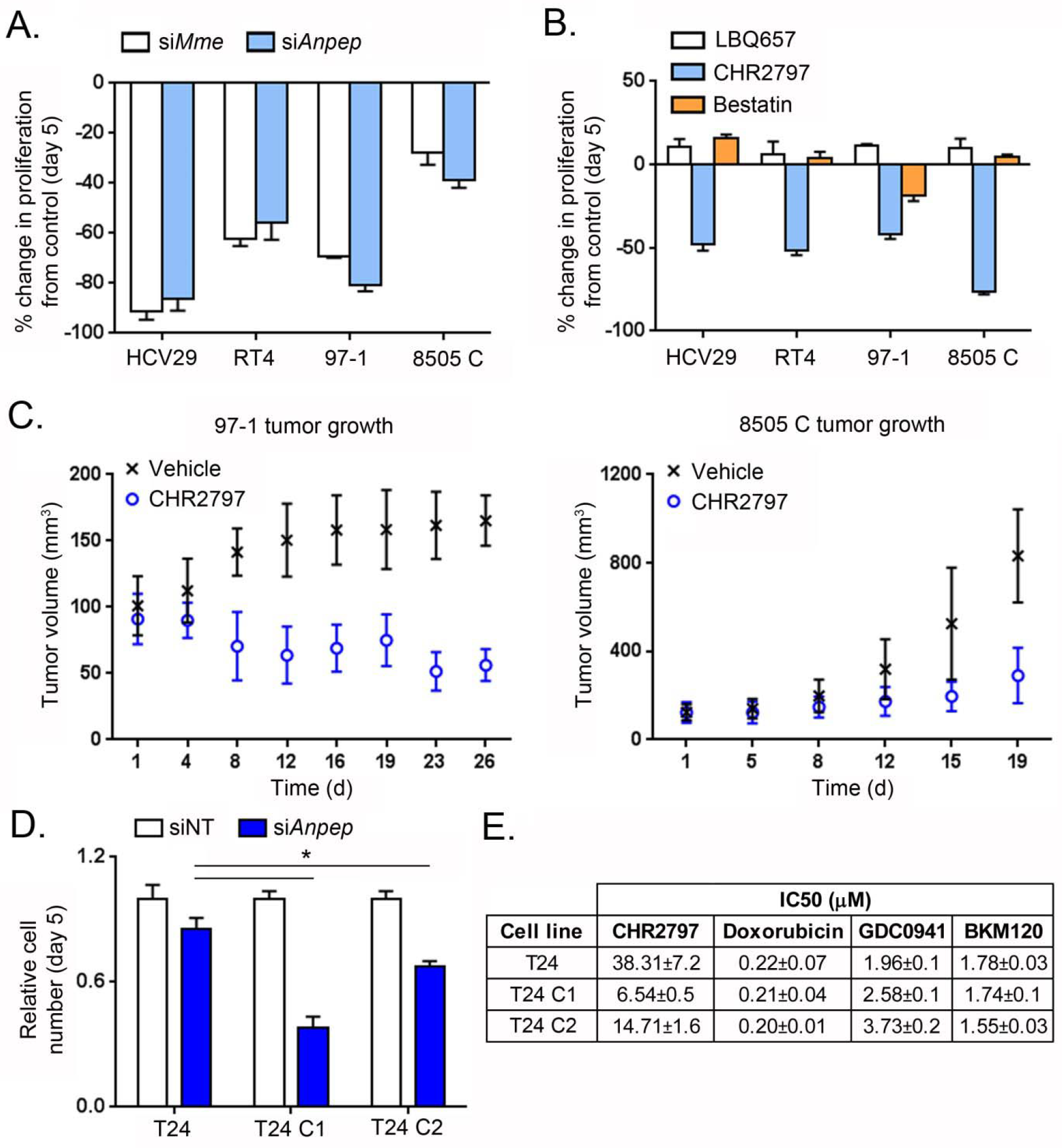
Functional studies reveal a dependence on NEP and APN for the proliferation and survival of TSC1 null cancer models.

**A.** Proliferation studies with TSC1 mutant human bladder cancer and thyroid cancer cell lines show that all cell lines are sensitive to siRNA targeted to *Mme* and *Anpep*. Data were collected at day 5 post transfection using the Cell TitreGlo assay, and expressed as a percent change in luminescence from control (non-targeting siRNA). **B.** Proliferation studies with TSC1 mutant human bladder cancer and thyroid cancer cell lines show cells sensitive to a biochemical inhibitor of APN, but not NEP. Treatment with 20 μM LBQ657, a biochemical inhibitor of NEP, did not inhibit cellular proliferation compared to vehicle treated cells. Treatment with 20 μM bestatin or CHR2797, two biochemical inhibitor of APN, showed that bestatin had limited impact of cellular proliferation, while CHR2797 potently inhibited proliferation. CHR2797 also inhibited proliferation in a dose dependent fashion at 2 and 20 μM. Data are expressed as a percentage change in luminescence signal compared to vehicle treated cells, and cell number was acquired with the Cell TiterGlo assay after drug exposure for 5 days. **C.** Treatment of nu/nu mice bearing subcutaneous 97-1 (left) or 8505 C (right) xenografts with 100 mg/kg CHR2797 suppressed tumor growth over 20-26 days in vivo compared to vehicle treated mice (200 μL of saline per mice per day). Treatments were administered once daily via i.p. injection. **D.** Proliferation studies show that suppression of APN with siRNA results in stronger antiproliferative effects in T24 C1 and T24 C2 compared to parental T24. Data were collected at day 5 post transfection using the Cell TitreGlo assay. \**P*<0.01 **E.** A grid summarizing the IC50 values acquired for CHR2797, doxorubicin, GDC0941, and BKM120 shows in parental T24 and the T24 sublines with CRISPR knockdown. T24 C1 and C2 are more sensitive to CHR2797 compared to parental T24. By comparison, all cell lines were equally sensitive to doxorubicin and PI3K inhibitors. Data were collected at day 5 post treatment using the Cell TitreGlo assay.

We next determined if loss of TSC1 sensitizes cells to ablation or inhibition of APN in vitro. Genetic silencing of *Anpep* mRNA via siRNA resulted in a statistically larger antiproliferative effect in T24 C1, and T24 C2 compared to parental T24 (**Figure 2D** and **Supplemental Figure 11**). Dosing CHR2797 revealed that the IC_50_ for growth inhibition in T24 C1 and C2 knockout cells was ~3-6 fold lower than parental T24 (**Figure 2E** and **Supplemental Figure 12**). By comparison, T24, T24 C1, and T24 C2 were equally sensitive to doxorubicin, BKM120 (pan p110 inhibitor), and GDC0941 (p110α isoform inhibitor).

CHR2797 was previously shown to suppress mTORC1 signaling, and this struck us as a possible mechanism for the sensitivity of TSC1 null cell lines.^24^ However, treating 97-1 cells with CHR2797 had no impact on the phosphorylation levels of mTORC1 substrates in our hands (**Figure 3A**). Moreover, treatment of TSC1 null cell lines with RAD001, BEZ235, or INK128 did not reduce APN expression, suggesting that the induction of APN by loss of TSC1 is mTORC1 independent (**Figure 3B** and **Supplemental Figure 13A** and **13B**). Consistent with this model, APN expression levels were also not upregulated in *Tsc2*^-/-^ versus *Tsc2*^+/+^ MEFs (nor was APN expression detectable in *Tsc2*^+/+^, see **Figure 3C** and **Supplemental Figure 14**).

**Figure 3.**
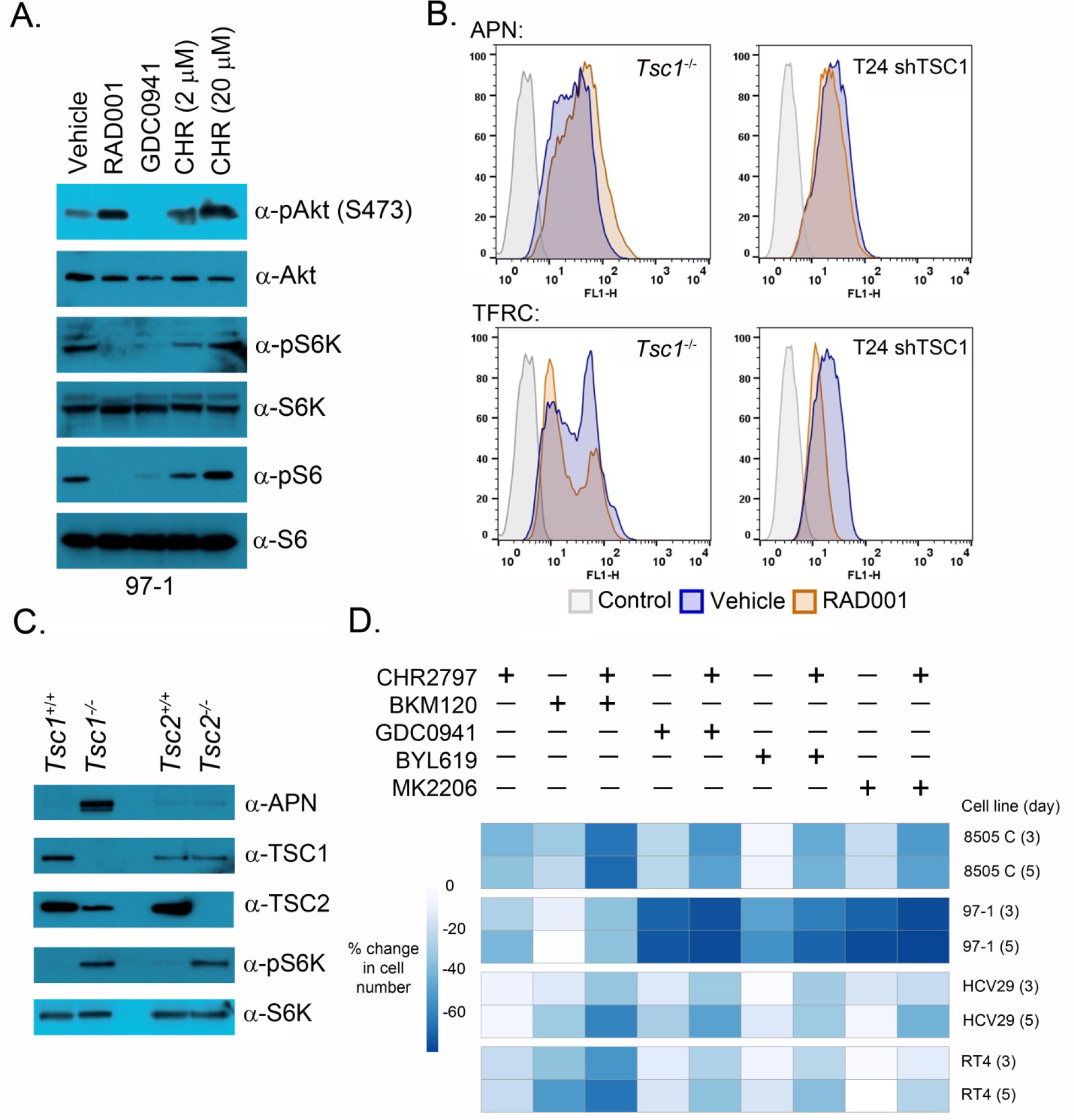
Mechanistic and pharmacological studies show that greater antiproliferative effects can be achieved by combining CHR2797 treatment with human ready inhibitors of the PI3K/Akt/mTOR signaling axis.

**A.** Immunoblot data showing that CHR2797 does not suppress mTORC1 signaling in vitro. 97-1 cells were exposed to the indicated dose of CHR2797, RAD001, or GDC0941 for 4 hours prior to lysing the cells for immunoblot. **B.** Representative histograms showing that RAD001 treatment in *Tsc1*^-/-^ MEFs and T24 C2 does not alter the expression of APN. TFRC expression changes were evaluated in parallel treatment arms, and RAD001 suppressed TFRC expression levels in each cell line, as expected. **C.** Immunoblot data showing that APN expression is not expressed in *Tsc2*^+/+^ or *Tsc2*^-/-^ MEFs. **D.** A heat map representing cellular proliferation data showing that TSC1 mutant human bladder and thyroid cancer cell lines are more sensitive to combination treatment with CHR2797 and inhibitors of the PI3K/Akt/mTOR signaling axis. Cells were exposed to the indicated drug and concentration for three or five days prior to evaluating cell number with Cell TitreGlo. The data were normalized to the luminescence signal associated with vehicle treated cells.

The observation that induction of APN by loss of TSC1 in a mTORC1 independent manner led us to propose that combined APN and mTORC1 inhibition may result in greater antiproliferative effects in TSC1 mutant cells. To this end, we treated TSC1 null cancer cell lines with CHR2797 alone and in combination with BKM120, GDC0941, BYL719 (p110α isoform inhibitor), or MKK2206 (pan Akt inhibitor). Proliferation of all cell lines was sensitive to kinase inhibition with single agents at 3 or 5 days post treatment but proliferation was significantly slower when co-treat with CHR2797 (**Figure 3D** and **Supplemental Figure 15**). This suggest that while APN and mTORC1 are both induced through the loss of TSC1, they act through independent mechanism in driving proliferation.

## Discussion

The overall goal of this manuscript was to develop new strategies to treat clinically problematic cells arising from the loss of the prominent tumor suppressor complex TSC1/TSC2. An unbiased proteomics screen in *Tsc1*^-/-^ and *Tsc1*^+/+^ MEFs identified 12 candidate cell surface antigens, five of which were strongly induced by TSC1 loss. Deeper study of the two most upregulated antigens, NEP and APN, suggest they may be potential drug targets, as silencing either protease potently inhibited the proliferation of TSC1 mutant human cancer cells, while treatment with the human ready APN inhibitor CHR2797 suppressed proliferation in vitro and in vivo. While human TSC1 knockout cells appeared to be more sensitive to genetic and biochemical suppression of APN compared to an isogenic parental cell line, we were also struck by the observation that neither NEP nor APN expression was dependent on TSC2 or mTORC1. This observation led to the non-obvious insight that the bioactivity of APN inhibition can be augmented combined suppression of mTORC1 signaling.

To our knowledge, no prior association has been established between TSC1 loss, and NEP or APN overexpression, though NEP and APN expression has been observed in human cancers with frequent TSC1 mutation^25–29^. TSC1 loss may upregulate either protein via a transcriptional mechanism, as mRNA levels are higher in *Tsc1*^-/-^ versus *Tsc1*^+/+^ MEFs. Prior research showing that TSC1 can transduce TGFβ signaling to Smad2 and Smad3 independent of TSC2 and mTORC1 may provide an important mechanistic lead^30^, and we are currently working to elaborate the functional connection between TSC1, NEP and APN more thoroughly.

Fully exploiting the enhanced sensitivity of TSC1 mutant cancer cells to APN inhibition will likely require more selective APN modulators. Typical of small molecule APN inhibitors, CHR2797 harbors some activity against other M1 aminopeptidases, including leucine aminopeptidase and puromycin-sensitive aminopeptidase^24,31^. That said, our antitumor data with CHR2797 may have immediate clinical impact, as this drug is generally well tolerated in humans up to doses of 240 mg/d^32,33^, and radiographic responses have been reported for doses proportionate to those used for our animal studies. We are also hopeful that our data will motivate interest in combining CHR2797 with any of the available clinically active PI3K pathway inhibitors, perhaps in setting like metastatic bladder cancer, given the frequency of TSC1 mutation and the standing interest in treating this malignancy with mTORC1 inhibitors.

Although NEP and APN emerged as obvious targets for immediate study, some of the other hits emerging from the proteomics screen may have utility. For instance, TFRC is a well-established target for molecular imaging^34–36^, and radiolabeled transferrin molecules may be useful for detecting cells with TSC1 mutations. To our knowledge, GSGL1 and AGRL2 are essentially unstudied as drug or diagnostic targets. The G-coupled protein receptor AGRL2 in particular may emerge as an interesting new target, owing to its limited expression in normal tissues. More generally, our success in identifying new drug targets for TSC1 and RAS driven cancers using cell surface proteomics argues strongly that this approach may scale well to other common oncogenic drivers^37^.

## Materials and Methods

### General methods

*Tsc1*^-/-^, *Tsc1*^+/+^, *Tsc2*^-/-^, and *Tsc2*^+/+^ MEFs were kindly provided by Professor David Kwiatkowski, and were maintained in high-glucose and glutamine containing DMEM supplemented with 10% FBS and 1% penicillin-streptomycin. The human bladder cancer cell lines T24, 5637, RT4, HCV29, and TCCSUP were purchased from ATCC and subcultured according to the manufacturer’s recommendations. The human thyroid cancer cell line 8505 C was purchased from Sigma-Aldrich and subcultured according to the manufacturer’s recommendations. The bladder cancer cell line 97-1 was kindly provided by Professor Margaret Knowles, and subcultured in Hams F12 supplemented with 1% FBS, 1x penicillin-streptomycin, 1x insulin-transferrin-selenium, 2 mM glutamine, 1x NEAA and 1μg/ml hydrocortisone. LBQ657 was purchased from Cayman Chemical. Bestatin was purchased from Sigma-Aldrich. CHR2797 was purchased from MedKoo Biosciences. GDC0941 was purchased from LC Laboratories. BKM120 was purchased from AdipoGen. Doxorubicin, MK-2206 and BYL-719 were purchased from Adooq Biosciences. All small molecule drugs were used without further purification. Antibodies to total Akt, p-Akt (S473), total S6, p-S6 (S235/236), total S6K, p-S6K (T389), APN (mouse, CD13), TSC1 and TSC2 were acquired from Cell Signaling Technologies and used at a 1:1000 dilution. Actin (clone AC-15) was purchased from Sigma Aldrich and used at a 1:5000 dilution. The NEP (CD10) antibody was purchased from Invitrogen and used at a 1:1000 dilution. The antibody to APN (human, CD13) was purchased from Proteintech and used at a 1:1000 dilution. Primary antibodies and their respective sources for flow cytometry are listed in **Supplemental Table 2**. Primers for rtPCR were synthesized by Integrated DNA Technologies, and a full list of primer sequences appear in **Supplemental Table 3**.

### SILAC, membrane protein enrichment, and LC-MS/MS

*Tsc1*^+/+^ and *Tsc1*^-/-^ MEFs were cultured in DMEM SILAC media (Thermo Fisher Scientific) containing L-[^13^C_6_,^15^N_2_]lysine and L-[^13^C_6_,^15^N_4_] arginine (heavy label) (Thermo Fisher Scientific) or L-[^12^C_6_,^14^N_2_]lysine and L-[^12^C_6_,^14^N_4_]arginine (light label) for 5 passages to ensure full incorporation of the isotope labeling on cells. Cells were grown to 70% confluence and harvested after 6 hours of serum starvation. Cells were mixed at a 1:1 cell count ratio prior to cell surface capture enrichment. Briefly, live cells were treated with a sodium periodate buffer (2 mM NaIO_4_, PBS pH 6.5) at 4°C for 20 mins to oxidize terminal sialic acids of glycoproteins. Aldehydes generated by periodate oxidation were then reacted with biocytin hydrazide in a labeling buffer (1 mM biocytin hydrazide (biotium), 10 mM analine (Sigma), PBS pH 6.5) at 4°C for 90 mins. Cells were then washed four times in PBS pH 6.5 to remove excess biocytin-hydrazide and flash frozen.

Frozen cell pellets were lysed using RIPA buffer (VWR) with protease inhibitor cocktail (Sigma-Aldrich; St. Louis, MO) at 4°C for 30 mins. Cell lysate was then sonicated, clarified, and incubated with 500μL of neutravidin agarose slurry (Thermo Fisher Scientific) at 4°C for 30 mins. The neutravidin beads were then extensively washed with RIPA buffer, high salt buffer (1M NaCl, PBS pH 7.5), and urea buffer (2M urea, 50mM ammonium bicarbonate) to remove non-specific proteins. Samples were then reduced on-bead with 5mM TCEP at 55°C for 30 mins and alkylated with 10mM iodoacetamide at room temperature for 30 mins. To release bound proteins, we first performed an on-bead digestion using 20μg trypsin (Promega; Madison, WI) at room temperature overnight to remove any non-specific protein binders. The neutravidin beads were extensively washed again with RIPA buffer, high salt buffer (1M NaCl, PBS pH 7.5), and urea buffer (2M urea, 50mM ammonium bicarbonate). To release trypsin digested N-glycosylated peptides bound to the neutravidin beads, peptides were released using 2500U PNGase F (New England Biolabs; Ipswich, MA) at 37°C for 3 hours and eluted using a spin column. PNGase F released peptides were then desalted using SOLA HRP SPE column (Thermo Fisher Scientific) using standard protocol, dried, and dissolved in 0.1% formic acid, 2% acetonitrile prior to LC-MS/MS analysis.

### Mass spectrometry analysis

Approximately 1μg of peptide was injected to a pre-packed 0.75mm x 150mm Acclaimed Pepmap C18 reversed phase column (2μm pore size, Thermo Fisher Scientific) attached to a Q Exactive Plus (Thermo Fisher Scientific) mass spectrometer. The peptides were separated using the linear gradient of 3-35% solvent B (Solvent A: 0.1% formic acid, solvent B: 80% acetonitrile, 0.1% formic acid) over 120 mins at 300μL/min. Data were collected in data-dependent mode using a top 20 method with dynamic exclusion of 35 secs and a charge exclusion setting that only sample peptides with a charge of 2, 3, or 4. Full (ms1) scans spectrums were collected as profile data with a resolution of 140,000 (at 200 *m*/*z*), AGC target of 3E6, maximum injection time of 120 ms, and scan range of 400 − 1800 *m*/*z*. MS-MS scans were collected as centroid data with a resolution of 17,500 (at 200 *m*/*z*), AGC target of 5E4, maximum injection time of 60 ms with normalized collision energy at 27, and an isolation window of 1.5 *m*/*z* with an isolation offset of 0.5 *m*/*z*.

### Proteomics data processing

Peptide search for each individual dataset was performed using ProteinProspector (v5.13.2) against 16916 mouse proteins (Swiss-prot database, obtained March 5, 2015) with a false discovery rate (FDR) of <1%. Quantitative data analysis was performed using Skyline (University of Washington) software using the ms1 filtering function. Specifically, spectral libraries from forward and reverse SILAC experiments were analyzed together such that ms1 peaks without an explicit peptide ID would be quantified based on aligned peptide retention time. An isotope dot product of at least 0.8 was used to filter out low quality peptide quantification, and a custom report from skyline was then exported for further processing and analysis using R: A language and environment for statistical computing. To ensure stringent quantification of the surface proteome, only peptides with N to D deamidation modification were included. Forward and reverse SILAC datasets were then combined and reported as median log_2_ enrichment values for *Tsc1*^-/-^ MEFs. For Gene Set Enrichment Analysis (GSEA), genes were ranked by median log2 enrichment values and analyzed against a curated mouse version of the MSigDB (http://bioinf.wehi.edu.au/software/MSigDB/) using the fast pre-ranked gene set enrichment analysis (fgsea) package from Bioconductor.

### Flow cytometry

All cell lines were grown in T75 flasks. Cells were washed with phosphate-buffered saline (PBS) and detached from cell culture dishes by 0.04% EDTA in PBS solution, centrifuged and washed with PBS again. Then the cells were fixed by 1% formaldehyde in PBS solution at 4 °C overnight. The cells were washed centrifuged and washed with PBS, and then counted. Cells were re-suspended in 3% BSA in PBS solution to a concentration of 0.7 million cells/100 μl. The corresponding primary antibodies were added based on the vendor’s recommendations, otherwise, 5 μl antibodies were added and incubated at room temperature for 30 min. Cells were washed three times with 3% BSA in PBS solution and re-suspended in 200 μl 3% BSA in PBS solution. 1 μl corresponding secondary antibodies were added and incubated at room temperature for 30 min if the primary antibodies were unconjugated. Cells were washed three times with 3% BSA in PBS solution and re-suspended in 400 μl PBS. Cells were analyzed on BD FACS Calibur flow cytometer.

### Immunoblot

Cell pellets were lysed in RIPA buffer with protease and phosphatase inhibitor cocktails (Calbiochem) and then resolved using 1D SDS PAGE. Protein concentration was determined with a Bradford absorbance assay, and equal amounts of protein (10-30 μg of lysate) were separated by SDS-PAGE, transferred to PVDF membranes, and immunoblotted with specific primary and secondary antibodies. Immunoreactive bands were visualized using enhanced chemiluminescence (ECL) and detected by chemiluminescence with the ECL detection reagents from Thermo Scientific.

### Real time PCR

Cellular RNA was harvested with a RNAeasy mini kit (Qiagen) using a Qiashredder to disrupt cell pellets. The purity and concentration of RNA was quantified using a NanoDrop spectrometer (ThermoScientific), and 1.5 μg of RNA was converted to cDNA with a high capacity cDNA reverse transcription kit (Applied Biosystems). Relative changes in mRNA levels were assessed with a Pikoreal rtPCR cycler (Thermo Fisher Scientific). ΔCt was calculated using the respective actin control, and ΔΔCt was calculated by normalizing ΔCt values to that of vehicle control. Data were expressed as 2^−ΔΔCt^. All measurements were performed with at least 4 replicates, and are representative of at least two independent experiments.

### Cellular proliferation studies

Proliferation/viability of cells was detected by using the CellTiter-Glo Luminescent Cell Viability One Solution Assay (Promega, Madison, WI). For the drug treatment assay, cells (1×10^3^ cells for MEFs and 2×10^3^ cells for human cancer cell lines) were plated on 96-well plates and incubated overnight for cell attachment. Cells were then treated for designed days with different drug treatments. All measurements were performed with at least 4 replicates, and are representative of at least two independent experiments.

### Gene suppression with shRNA knockdown or CRISPR/Cas9 mediated genetic knockout

Five pLKO.1 vectors containing discrete shRNA sequences targeting TSC1 were purchased from the MISSION shRNA collection (Sigma Aldrich). The packaging plasmid pSPAX2 and the envelop plasmid pMD2.G were purchased from Addgene. To prepare lentiviral particles, HEK-293FT cells (1.5 x 10^6^) were plated, and after 18 hours, the medium was replaced with antibiotic free DMEM media containing 10% FBS. pLKO.1 (2 μg), pSPAX2 (1.8μg), and pMD2.G (0.6 μg) plasmids were mixed together in 70 μL Opti-MEM. In a separate Eppendorf, Lipofectamine 2000 (15 μL) was diluted in 70 μL Opti-MEM. The solutions containing plasmid and Lipofectamine were mixed together and added directly to a plate of HEK293 FT cells. After 12 h, the media was replaced with 5 mL of DMEM containing 1% pen/strep, 10% heat inactivated FBS, and 1.1% BSA. The virus containing media was harvested at 24 h, replaced with fresh media, and harvested again at 48 h. The virus was stored at 4° C or used immediately after filtration through a low-binding protein membrane (0.45 μm). For infection, 0.2 M bladder cancer cell lines were seed into the 10 cm^2^ plates overnight for attaching. Then the media was changed to fresh media containing 8 μg/mL of polybrene. After 2 h, 1 mL of virus media was added. After 24h, the media was replaced with complete media which contained 1 ug/mL puromycin. After selection by puromycin for two weeks, the knockdown effects were checked by western blot and the cells were stored for future studies. Of the five shRNA sequences assayed, we determined that optimal knockdown was achieved with the following targeting sequence: 5’

CCGGGCACTCTTTCATCGCCTTTATCTCGAGATAAAGGCGATGAAAGAGTGCTTTTTG-3’, clone ID: NM_000368.3-790s21c1, MISSION TRC no: TRCN0000327868. To achieve CRISPR mediated gene knockout of TSC1, a similar workflow was followed using the same packaging and envelope plasmids. Two targeting sequences were used in the lentiCRISPRv2 plasmid (Addgene): 5’-TCAGGCACCATGATGACAGA-3’ (T24 C1) and 5’-CGAGAGGAT GGATAAAC GAG-3’ (T24 C2)^38^. The targeting sequences were designed with CHOPCHOP software^39^. After puromycin selection of infected T24 cells for two weeks, single cell clones were isolated collected using FACS (BD FACSAria2) into a 96 well plate and allowed to expand in complete media prior to confirmation of TSC1 knockout via immunoblot.

### siRNA Transfection

ONTARGET Plus siRNA SMARTpools against *Mme* and *Anpep* were purchased from Dharmacon. A non-targeting SMARTPool siRNA cocktail was used as a negative control. Cells were transfected by Lipofectamine^®^ RNAiMAX Transfection Reagent according to the manufacturer’s protocol (Thermo Scientific). In brief, for a 6-well plate, 80,000 cells (in 2 mL medium) were plated in each well and incubated overnight for cell attachment. siRNA solutions were prepared Opti-MEM medium. After adding siRNA, the final concentration of siRNA was 40 nM in each well. 72 h after transfection, cells were counted by CellTiter-Glo. mRNA silencing was confirmed in separate wells using rtPCR.

### Apoptosis assay

Apoptosis of cells was detected by using the Caspase-Glo 9 assay (Promega). For detection of caspase activity and apoptosis, cells (2 x 10^3^/well) were plated on 96-well plates and incubated overnight for cell attachment. Cells were then treated with CHR2797 (final concentration: 2 μM) for 3 or 5 days. The final numbers were normalized by cell numbers. All measurements were performed with at least 4 replicates and were independently reproduced at least two times.

### Biochemical assays

Cellular APN activity was measured with a fluorometric aminopeptidase N activity assay kit (BioVision). Viable, intact cells (5 x 10^5^) were dispersed in 90 μL of assay buffer, and the reaction was initiated by the addition of 10X APN Substrate solution (10 μL). Substrate hydrolysis by APN was measured at Ex/Em = 384/502 nm every 30 sec for 20-45 min at 37°C. A similar workflow was applied to measure cellular NEP activity using a fluorogenic neprilysin activity assay kit (BioVision) using 1 x 10^5^ cells, the corresponding NEP substrate, and Ex/Em = 330/430 nm. In both cases, the data were plotted using PRISM, and a linear regression was applied to calculate the slope, which represents a rate of substrate hydrolysis proportional to cell surface enzyme expression. The activity was assayed in triplicate, and independently reproduced.

### In vivo anti-tumor assessment

All animal studies were performed in accordance with IACUC protocols and were approved by the Laboratory Animal Research Center at UCSF. Intact male *nu/nu* nude mice (Charles River) received 97-1 or 8505 C cells (1×10^6^) in the flank in a 1:1 PBS:matrigel suspension (Corning Matrigel Matrix High Concentration). When tumor volume reached 70-110 mm^3^, mice were randomly grouped into arms receiving vehicle (saline) or CHR2797 (100 mg/kg). Intraperitoneal injection was applied and the dose per day for the treatment group was 100 mg/kg in 200 μL formulation (10% DMSO, 10% Tween 80, 80% saline solution). Tumor dimensions were measured with calipers, and the volume was calculated using the following formula: 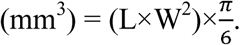

### Statistical analysis

Statistical evaluations were performed by Student’s t-test for paired data and by ANOVA for sets of data with multiple comparison points. Data were considered to be statistically significant if *P* < 0.05.

## Acknowledgements

The authors gratefully acknowledge Mr. Loc Huynh, Yung-hua Wang, and Tony Huynh for technical assistance, and Dr. Sarah Elmes for assistance with FACS. M.J.E. was supported by the 2013 David H. Koch Young Investigator Award from the Prostate Cancer Foundation and the American Cancer Society (130635-RSG-17-005-01-CCE). C.T. was supported by a postdoctoral fellowship from the Department of Defense Prostate Cancer Research Program (PC151060) Research from UCSF reported in this publication was supported in part by the National Cancer Institute of the National Institutes of Health under Award Number P30CA082103. The content is solely the responsibility of the authors and does not necessarily represent the official views of the National Institutes of Health.

